# NOTUM-MEDIATED WNT SILENCING DRIVES EXTRAVILLOUS TROPHOBLAST CELL LINEAGE DEVELOPMENT

**DOI:** 10.1101/2024.02.13.579974

**Authors:** Vinay Shukla, Ayelen Moreno-Irusta, Kaela M. Varberg, Marija Kuna, Khursheed Iqbal, Anna M. Galligos, John D. Aplin, Ruhul H. Choudhury, Hiroaki Okae, Takahiro Arima, Michael J. Soares

## Abstract

Trophoblast stem (**TS**) cells have the unique capacity to differentiate into specialized cell types, including extravillous trophoblast (**EVT**) cells. EVT cells invade into and transform the uterus where they act to remodel the vasculature facilitating the redirection of maternal nutrients to the developing fetus. Disruptions in EVT cell development and function are at the core of pregnancy-related disease. WNT-activated signal transduction is a conserved regulator of morphogenesis of many organ systems, including the placenta. In human TS cells, activation of canonical WNT signaling is critical for maintenance of the TS cell stem state and its downregulation accompanies EVT cell differentiation. We show that aberrant WNT signaling undermines EVT cell differentiation. Notum, palmitoleoyl-protein carboxylesterase (**NOTUM**), a negative regulator of canonical WNT signaling, was prominently expressed in first trimester EVT cells developing in situ and upregulated in EVT cells derived from human TS cells. Furthermore, NOTUM was required for optimal human TS cell differentiation to EVT cells. Activation of NOTUM in EVT cells is driven, at least in part, by endothelial PAS domain 1 (also called hypoxia-inducible factor 2 alpha). Collectively, our findings indicate that canonical WNT signaling is essential for maintenance of human trophoblast cell stemness and regulation of human TS cell differentiation. Downregulation of canonical WNT signaling via the actions of NOTUM is required for optimal EVT cell differentiation.

**SIGNIFICANCE:** Extravillous trophoblast (**EVT**) cells play a critical role in transforming the uterine environment into a supportive organ facilitating embryonic/fetal development. Insufficient EVT cell-dependent uterine transformation can lead to obstetrical complications, including early pregnancy loss, preeclampsia, intrauterine growth restriction, and preterm birth. These complications carry a significant burden of morbidity and mortality for both the mother and the fetus. Notum, palmitoleoyl-protein carboxylesterase, a WNT signaling antagonist, is involved in promoting and maintaining EVT cell differentiation. This process is essential for the proper development of the placenta and is crucial for a healthy pregnancy.

## INTRODUCTION

Development of the human placenta is necessary for proper embryonic development and successful pregnancy outcome (**1**). The hemochorial placenta develops through tightly regulated expansion and differentiation of trophoblast stem (**TS**) cells (**2**). Human TS cells can differentiate into two specialized cell types: syncytiotrophoblast and extravillous trophoblast (**EVT**) cells (**3-5**). Syncytiotrophoblast regulates maternal and fetal homeostasis through hormone production and control of the transfer of nutrients between maternal and fetal compartments, whereas EVT cells facilitate transformation of the uterus into a structure supportive of fetal growth and development (**4, 5**). EVT cells restructure uterine vasculature to optimize nutrient flow to the placenta (**6-8**). Abnormalities in EVT cell invasion and uterine spiral artery remodeling have been detected in an assortment of pregnancy disorders such as early pregnancy loss, preeclampsia, intrauterine growth restriction, and preterm birth (**9, 10**). Thus, experimental investigation of mechanisms underlying the derivation of the invasive EVT cell lineage could represent a key to understanding the etiology of placental dysfunction leading to pregnancy related disorders. At this juncture, there is a paucity of knowledge regarding the regulation of TS cells and especially their differentiation into EVT cells.

WNT signaling has been implicated as a key regulator of morphogenesis of many developing organ systems (**11**). WNT proteins represent a family of ligands that act by interacting with members of the Frizzled (**FZD**) receptor protein family (**11, 12**). Effective recognition by FZD proteins requires the post-translational addition of a palmitoleate moiety to WNT (**13, 14**), which is performed by Porcupine O-acyltransferase (**15**). Palmitoylated WNT is a target of Notum, Palmitoleoyl-protein Carboxylesterase (**NOTUM**), a secreted deacylase (**16, 17**). NOTUM removes the palmitoleate moiety from WNT, which destabilizes WNT interactions with FZD and negates downstream WNT/FZD-mediated signal transduction (**16, 17**). Canonical WNT signaling involves the redistribution of β-catenin (**CTNNB1**) from the cell membrane to the nucleus where it interacts with members of the lymphoid enhancer binding factor-1 (**LEF1**)/T cell factor (**TCF**) family of transcription factors (**12, 18**).

WNT signaling has been implicated in the regulation of placental development (**19-21**). WNT ligands are expressed within the early embryo (**22, 23**) and throughout the placentation site (**24**). A broad spectrum of activities has been ascribed to canonical WNT signaling in placental morphogenesis. These include roles in maintaining trophoblast cells in the stem/proliferative state (**25, 26**) and in promoting differentiation into syncytiotrophoblast (**27, 28**) and EVT cells (**29-31**). Supporting both the stem state and cell differentiation would seem to be opposing efforts but may reflect actions that are developmental stage and context dependent. Disruptions in canonical WNT signaling have also been connected to diseases of placentation (**32, 33**)

In this report, we evaluate the role of canonical WNT signaling in the regulation of human TS cell differentiation to EVT cells. The capture and ex vivo propagation of human TS cells represented a breakthrough for investigating human trophoblast cell differentiation (**25**). These cells can be maintained in a stem state or induced to differentiate into syncytiotrophoblast or EVT cells. We demonstrate that canonical activation of WNT signaling can block EVT cell differentiation and discover a critical role for NOTUM in facilitating EVT cell differentiation through attenuating canonical WNT signaling.

## RESULTS

### WNT activation disrupts EVT cell differentiation

Canonical WNT signaling supports maintenance of human TS cells in the stem state (**25**). Initially, using the human TS cell model, we monitored canonical WNT signaling by assessing active CTNNB1 (dephosphorylated on Ser-37 and Thr-41) using immunofluorescence in X,X CT-27 human TS cells maintained in the stem state and following EVT cell differentiation. Active CTNNB1 was abundant in stem cells but not in EVT cells, suggesting that WNT signaling was downregulated during EVT cell development (**Fig. 1A**) and supporting the involvement of WNT signaling in maintenance of the stem state. Standard culture conditions used to promote the TS cell stem state include the glycogen synthase kinase 3 (**GSK3**) inhibitor, CHIR99021, an activator of canonical WNT signaling (**25**). In contrast, culture conditions promoting EVT cell differentiation exclude CHIR99021 (**25**). We next sought to directly determine the consequences of activation of WNT signaling (CHIR99021) on EVT cell differentiation.

**Fig. 1.**
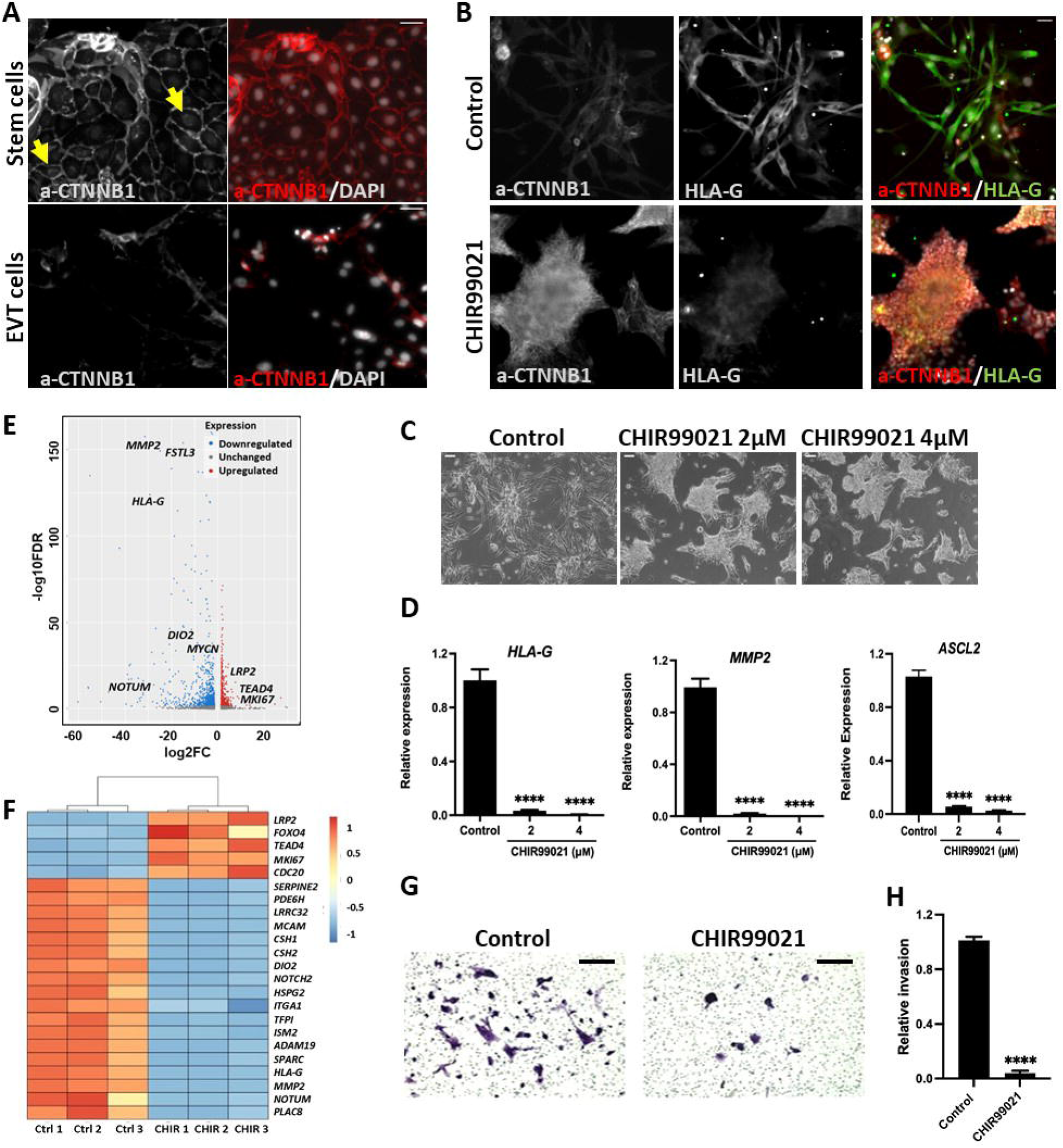
WNT activation disrupts extravillous trophoblast (**EVT**) cell differentiation. Experiments were performed with X,X CT-27 cells. *(A)* Immunocytochemistry of active CTNNB1 (a-CTNNB1) protein expression in stem and EVT cells. (Scale bar: 50 μm). 4’,6-Diamidino-2-phnylindole (**DAPI**) stains nuclei grey (right panel). Yellow arrows indicate active CTTNB1 in nuclei of TS cells. *(B)* a-CTNNB1 and HLA-G protein expression assessed in the presence of the WNT activator, CHIR99021(2 μM). DAPI nuclei are shown in grey (right panel). Merged fluorescence images of a-CTNNB1, HLA-G and DAPI images (Scale bar: 50 μm). *(C)* Phase contrast images depicting cell morphology of EVT cells in control conditions or in the presence of CHIR99021 on day 8 of EVT cell differentiation (Scale bar: 100 μm). *(D)* Transcript levels for EVT cell associated transcripts: HLA-G, MMP2 and ASCL2 on day 8 of EVT cell differentiation in control conditions or in the presence of CHIR99021, n=3, Graphs represent mean values ± standard error of the mean (**SEM**), one-way analysis of variance, Tukey’s post hoc test, ****P<0.0001. *(E and F)* A volcano plot and a heat map depicting RNA-sequencing analysis of control and CHIR99021 treated EVT cells (n=3 per group). *(E)* Blue dots represent significantly downregulated transcripts (P≤0.05) and a logarithm of base two-fold change of less than or equal to −2. Red dots represent significantly upregulated transcripts with P≤0.05 and a logarithm of base two-fold change of ≥2. *(F)* Heat map colors represent z-scores of reads per kilobase per million (**RPKM**) values. *(G)* Matrigel invasion assay of control or CHIR9902 (2 μM) treated cells under conditions promoting EVT cell differentiation. *(H)* The relative number of invading cells for control and CHIR9902 (2 μM) treated cells. Mean values ± SEM are presented, n=3, unpaired t-test, ****P<0.0001.

CHIR99021 was added as a supplement to the standard EVT cell differentiation culture conditions of X,X CT27 human TS cells (**25**). We examined the effects of WNT activation throughout EVT cell differentiation. Microscopy and immunofluorescence analyses revealed that addition of CHIR99021 resulted in the accumulation of active CTNNB1 (**Fig. 1B**) and an inhibition of EVT cell differentiation. The elongated/spindle-shaped cells characteristic of in vitro EVT cell differentiation were less abundant following induction of WNT signaling (**Fig. 1C**). EVT cell signature transcripts (e.g. *HLA-G*, *MMP2*, and *ASCL2*) and major histocompatibility complex, class I, G (**HLA-G**) protein were decreased in TS cells exposed to EVT cell differentiating conditions supplemented with CHIR99021 (**Fig. 1B, D**). We extended this analysis to an investigation of the effects of WNT activation on the EVT cell transcriptome (**Fig. 1E, F and Dataset 1**). RNA sequencing (**RNA-seq**) identified 1,580 differentially expressed genes (**DEGs**, false discovery rate, P<0.05), including 599 transcripts upregulated and 981 transcripts downregulated by exposure to CHIR99021. Stem state signature transcripts (e.g. *LRP2*, *TEAD4*, *MKI67*) were upregulated and EVT cell signature transcripts (e.g. *HLA-G*, *MMP2*, *ASCL2*, *FSTL3*, *NOTUM*) were downregulated (**Fig. 1F**). This differential gene expression pattern was validated by reverse transcriptase-quantitative PCR (**RT-qPCR**) (***SI Appendix,* Fig. S1**). In addition, trophoblast cell invasiveness was examined using Matrigel^Ò^ Transwell assays. WNT activation inhibited EVT cell migration (**Fig. 1G and H**). We next examined the short-term effects of CHIR99021 withdrawal during the TS cell stem state and the actions of CHIR99021 during an extended EVT cell differentiation culture protocol (days 6 to 14 of EVT cell differentiation). Evidence for EVT cell differentiation was observed within 48 h of CHIR99021 withdrawal as demonstrated by significant enhancement of the expression of some EVT signature transcripts (***SI Appendix,* Fig. S2**). WNT activation during extended EVT cell differentiation downregulated the expression of several EVT cell signature transcripts (*CDH5, MMP12, FLT4, SERPINE1, TIMP3, PAPPA2, ISM2, NOTUM, HLAG, MMP2, SNAI1*), except for *AOC1* expression, which was significantly stimulated with the high concentration of CHIR99021 (***SI Appendix,* Fig. S3**). *AOC1* expression is a measure of late-stage EVT cell maturation (**34**).

Overall, these results demonstrated that canonical WNT signaling had a prominent restraining role on the initiation of EVT cell differentiation and interfered with the maintenance of the EVT cell phenotype. We next sought evidence to determine whether canonical WNT signaling was operational in the first trimester human placenta.

### Canonical WNT signaling in the human placentation site

To assess the role of WNT signaling in the developing human placenta, we examined the distribution of active CTNNB1 and major histocompatibility complex, class I, G (**HLA-G**) protein in first trimester placental tissue specimens. Activated CTNNB1 was prominently expressed in cytotrophoblast populations, including basal cytotrophoblast situated at the base of the EVT cell column and cells within the transition zone of the EVT cell column (**Fig. 2A**). Co-expression of active CTNNB1 and HLA-G was not observed within the EVT cell column (**Fig. 2B and C; *SI Appendix,* Fig. S4**). We extended this analysis to the decidual bed and observed heterogeneity among HLA-G positive EVT cells (**Fig. 2G-I**). Some HLA-G positive EVT cells expressed active CTNNB1, while other HLA-G positive EVT cells lacked active CTNNB1 expression (**Fig. 2I**). Thus, EVT cells within the decidual bed can be distinguished based on their responsiveness to WNT signaling. At this juncture, human TS cells do not represent an effective model for investigating EVT cells expressing activated CTNNB1.

**Fig. 2.**
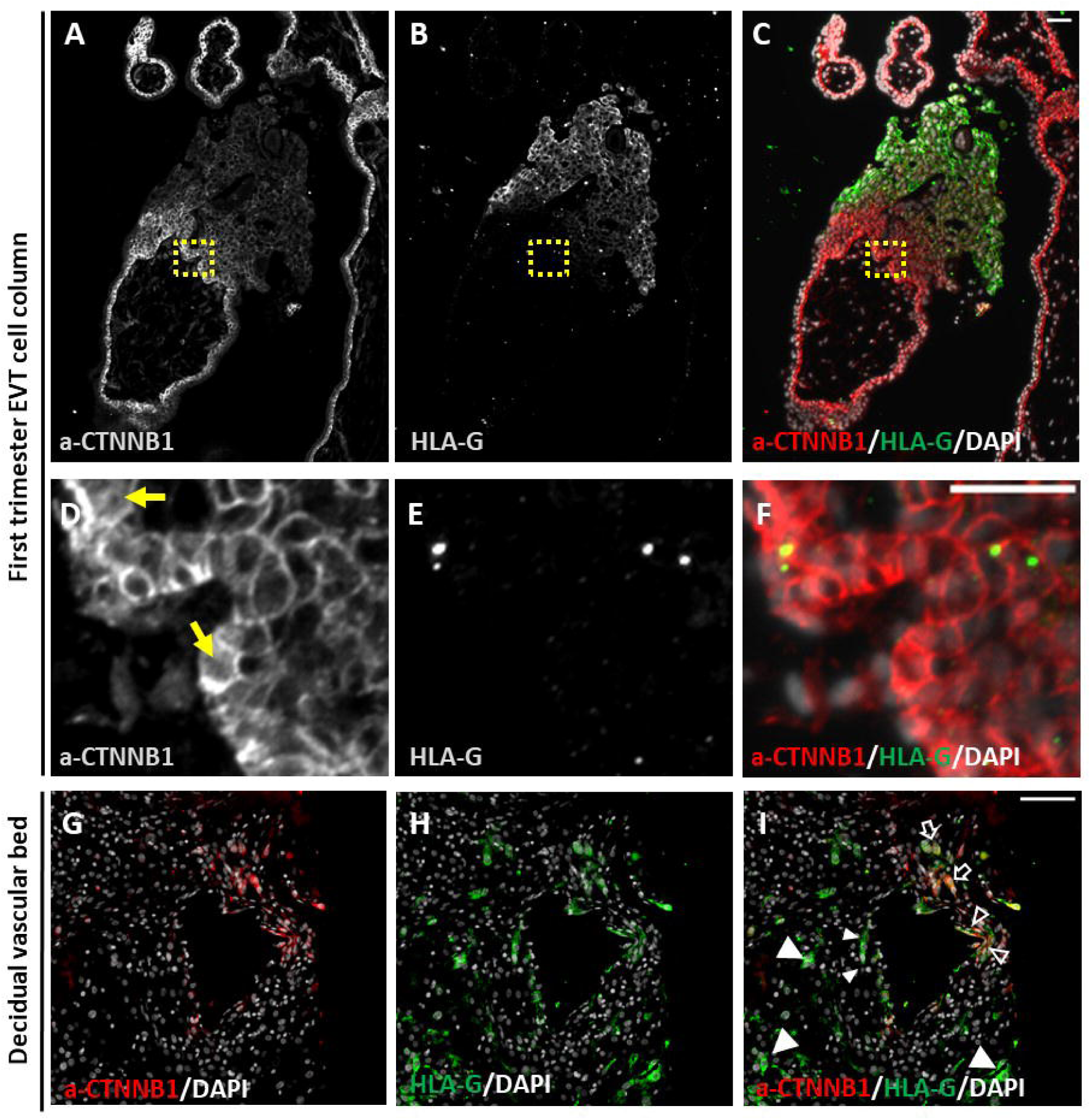
Immunofluorescence of first trimester human placenta at 12 weeks of gestation tested for the presence of active CTNNB1 (**a-CTNNB1**) and HLA-G. *(A)* a-CTNNB1 is expressed in cytotrophoblast, whereas *(B)* HLA-G is expressed in cells of the EVT cell column but not in cytotrophoblast or syncytiotrophoblast compartments. *(C)* Merged fluorescence images of a-CTNNB1 (red) and HLA-G (green). (D-F) High magnification images of yellow insets shown in panels A-C, respectfully. Yellow arrows indicate a-CTTNB1 in nuclei of cytotrophoblasts (Scale bar: 50 μm). Decidual bed tissue immunostained for a-CTNNB1 (red, panel G), HLA-G (green, panel H), and merged fluorescence images of a-CTNNB1 (red) and HLA-G (green,panel I). White arrowheads indicate EVT cells positive for HLA-G but negative for a-CTNNB1. Unfilled white arrowheads indicate EVT cells co-expressing HLA-G and a-CTNNB1. DAPI positive nuclei are shown in grey. Scale bar: 50 μm.

We were next directed to elucidating endogenous mechanisms during EVT cell differentiation responsible for liberating TS cells from the restraining actions of WNT signaling.

### NOTUM is expressed in EVT cells

Inspection of RNA-seq datasets from human TS cells in stem and EVT cell differentiated states (**25, 35**) showed that *NOTUM* was prominently expressed during EVT cell differentiation. The EVT cell-dependent upregulation of NOTUM was verified by RT-qPCR, western blotting, and immunofluorescence (**Fig. 3A-C**). NOTUM was co-localized with HLA-G in EVT cells (***SI Appendix,* Fig. S5)**. These observations prompted an investigation of *NOTUM* expression in the first trimester human placenta. *NOTUM*, *CDH1,* and *PLAC8* transcripts were localized by in situ hybridization. *CDH1* is a marker of a proliferative population of cytotrophoblast situated at the base of EVT cell columns (basal cytotrophoblast, **35, 36**), whereas *PLAC8* is distinctively expressed in EVT cells situated in the distal part of EVT cell columns (**35, 37**). Within the EVT cell column *NOTUM* did not co-localize with *CDH1* (**Fig. 3D**) but instead co-localized with *PLAC8* (**Fig. 3E**). This analysis was extended to decidual bed specimens possessing invasive EVT cells (**38**). NOTUM was co-localized to a subset of HLA-G positive EVT cells (**Fig. 3F**) but was not co-localized with activated CTNNB1(**Fig. 3G**). These findings are consistent with the detection of NOTUM expression in second trimester EVT cells (**39**) and the heterogeneity of HLA-G positive EVT cells (**Fig. 2I**). Thus, NOTUM is expressed in EVT cells differentiated from TS cells in vitro and within invading EVT cells identified in situ within first and second trimester human placentation sites.

**Fig. 3.**
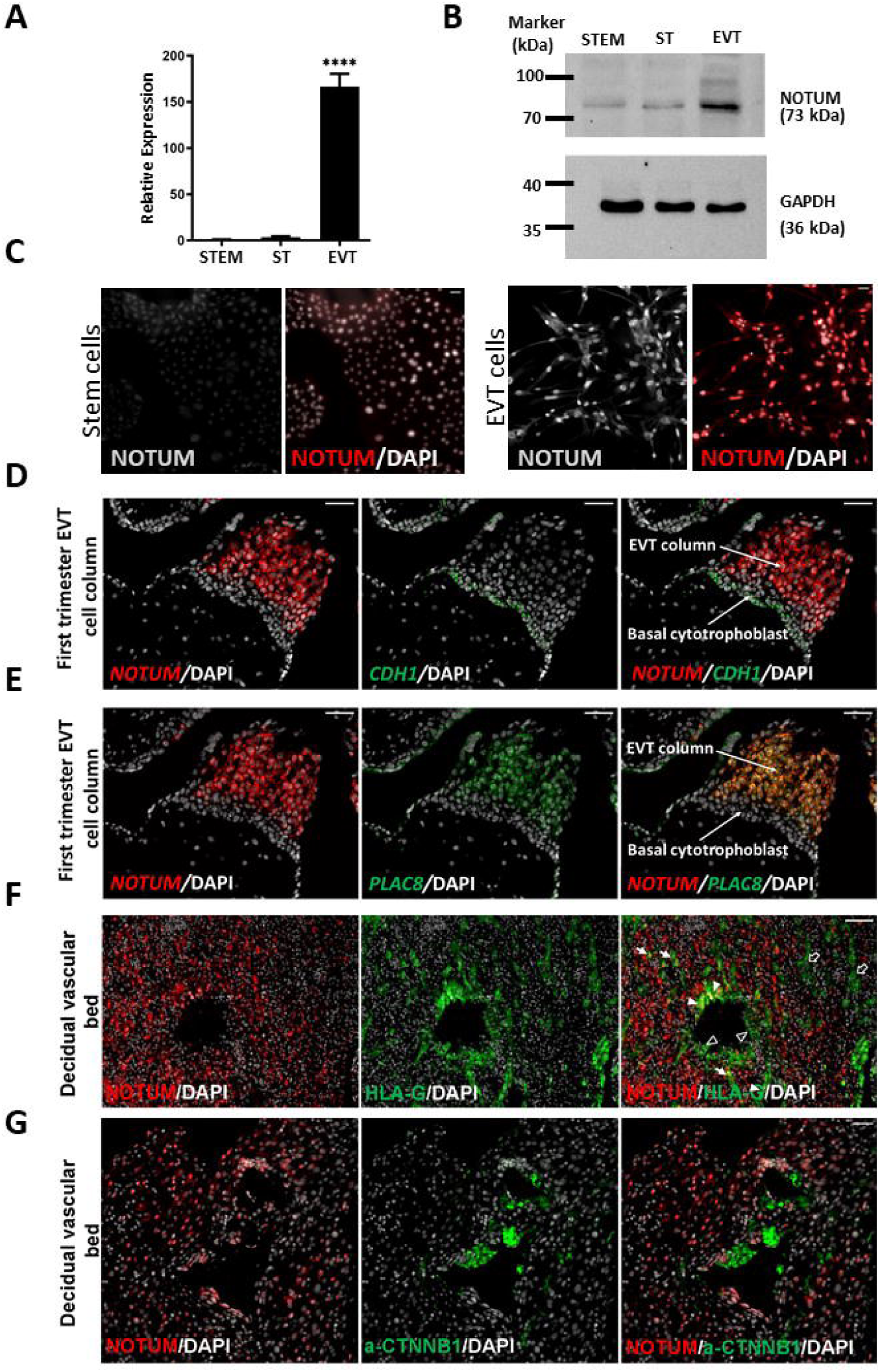
NOTUM is expressed in EVT cells. *(A)* Transcript levels of *NOTUM* in X,X CT-27 TS cells maintained in the stem state (**STEM**) or induced to differentiate into syncytiotrophoblast (**ST**) or day 8 EVT differentiation cells. Graphs represent mean values ± SEM, one-way analysis of variance, Tukey’s post hoc test, n=4, ****P<0.0001. *(B)* Western blot detection of NOTUM and GAPDH expression in X,X CT-27 TS cells maintained in the stem state or induced to differentiate into ST or EVT cells. *(C)* Immunofluorescence localization of NOTUM protein in TS cells maintained in the stem state or induced to differentiate into EVT cells. DAPI positive nuclei are shown in grey. *(D, E)* In situ hybridization of first trimester human placenta at 12 weeks of gestation probed for *NOTUM* (red), *CDH1* (green), or *PLAC8* (green). *NOTUM* (red) is expressed in EVT cell columns but not in basal cytotrophoblast. DAPI positive nuclei are shown in grey. *(F)* Decidual bed tissue immunostained for NOTUM (red) and HLA-G (green). White arrowheads indicate EVT cells positive for HLA-G and NOTUM. Unfilled white arrowheads indicate EVT cells positive for HLA-G and negative for NOTUM. (G) Decidual bed tissue immunostained for NOTUM (red) and a-CTNNB1 (green). DAPI positive nuclei are shown in grey. Scale bars: 50 μm.

### NOTUM promotes EVT cell differentiation

We hypothesized that NOTUM attenuates WNT-mediated inhibition of EVT cell differentiation. To test our hypothesis, we used a *loss-of-function* approach to determine whether NOTUM is required for EVT cell differentiation. NOTUM expression was silenced in X,X CT27 human TS cells using stable lentiviral-mediated delivery of short hairpin RNAs (**shRNA**) targeted to NOTUM. Disruption of NOTUM expression was verified by RT-qPCR and western blotting (**Fig. 4A, B**). TS cells expressing NOTUM shRNAs failed to show appropriate morphologic indices of EVT cell differentiation (**Fig. 4C**) and did not exhibit the characteristic upregulation of signature EVT cell transcripts (e.g., *HLA-G, MMP2, ASCL2, FSTL3, TFPI*) (**Fig. 4D**), or HLA-G protein expression (**Fig. 4E**). We extended this analysis to an investigation of the effects of NOTUM silencing on the EVT cell transcriptome (**Fig. 4F, G and Dataset 2**). RNA-seq analysis identified 542 DEGs (false discovery rate, P<0.05), including 195 upregulated transcripts and 347 downregulated transcripts in NOTUM knockdown cells. Consistent with morphologic observations and candidate transcript assessments, we identified a significant upregulation of stem state-specific transcripts (e.g., *LRP2*, *TEAD4*, *YAP1*, *F3*) and downregulation of EVT cell-specific transcripts (e.g., *HLA-G*, *MMP2*, *TFPI*, *ASCL2*, *CDKN1C*, *PLOD2*) (**Fig. 4G, Dataset 2, *SI Appendix,* Fig. S6**). Disruption of NOTUM also inhibited acquisition of the invasive properties of EVT cells (**Fig. 4H, I**) and resulted in activation of canonical WNT signaling (***SI Appendix,* Fig. S7**). Although, CHIR99021 exposure and disruption of NOTUM both led to inhibition of canonical indices of EVT cell differentiation, differences were noted in gene expression with each treatment (***SI Appendix*, Fig. S8; Dataset 3**). Similar to roles for canonical WNT activation and NOTUM silencing on X,X CT27 EVT cell differentiation, these experimental manipulations negatively impacted X,Y CT29 EVT cell differentiation (***SI Appendix,* Fig. S9**). We also examined the consequences of nullifying the actions of NOTUM using a small molecule inhibitor, LP922056 (**40**). LP922056 similarly interfered with EVT cell differentiation (***SI Appendix,* Fig. S10**). Thus, NOTUM promotes EVT cell differentiation.

**Fig. 4.**
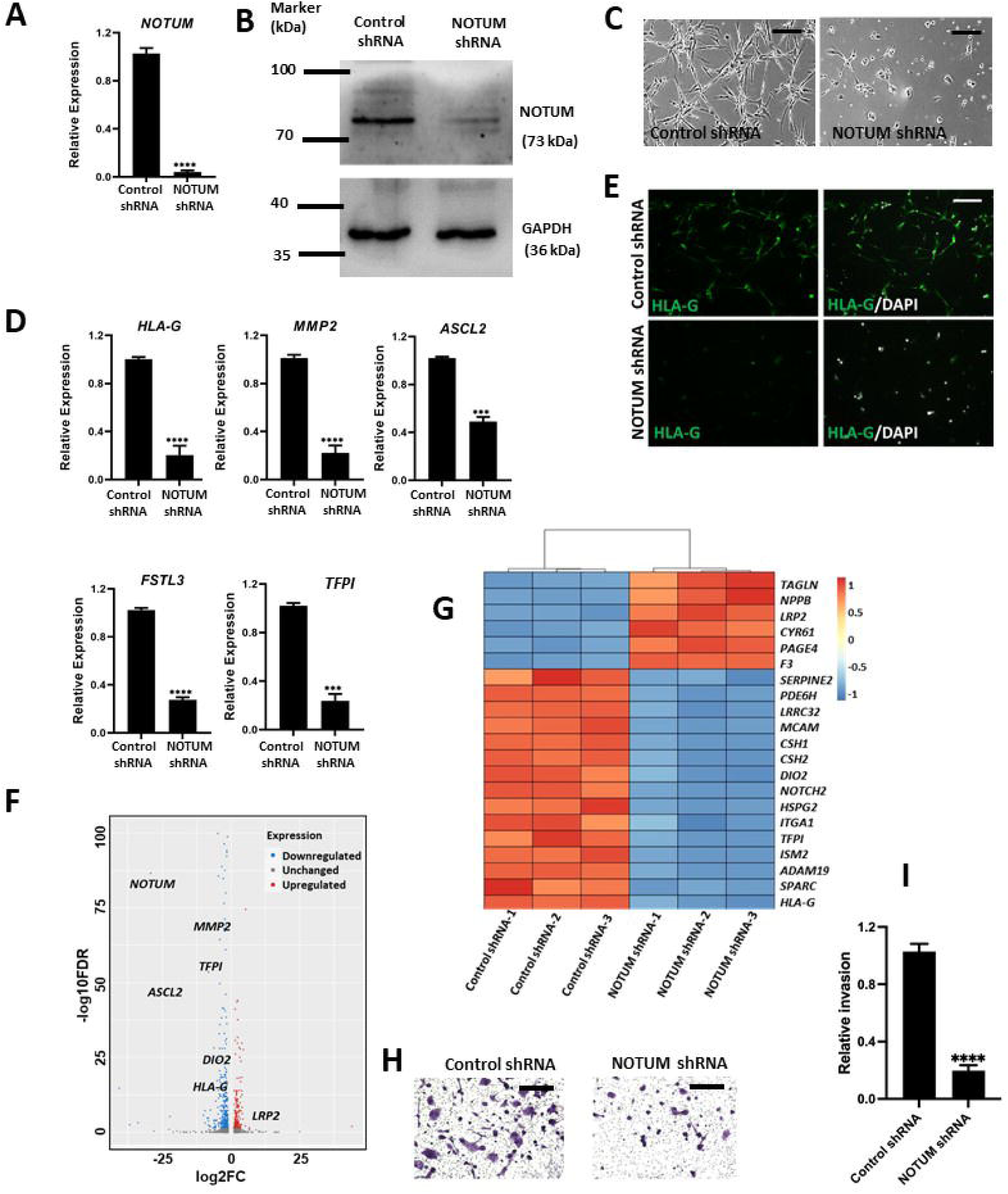
NOTUM regulates EVT cell differentiation. Experiments were performed with X,X CT-27 cells following 8 days of EVT cell differentiation. Effectiveness of NOTUM shRNA treatment was determined by *(A)* RT-qPCR and *(B)* western blotting. GAPDH was used as a loading control for the western blot. *(C)* Phase contrast images depicting cell morphology of TS cells cultured in conditions to promote EVT cell differentiation transduced with control shRNA or NOTUM shRNA (Scale bar: 200 μm). *(D)* Transcript levels of EVT cell markers (*HLA-G*, *MMP2*, *ASCL2*, *FSTL3*, and *TFPI*) in control shRNA and NOTUM shRNA treated cells following exposure to conditions that promote EVT cell differentiation. NOTUM knockdown decreased levels of transcripts characteristic of EVT cells (n=3 transductions), Graphs represent mean values ± SEM, n=3, unpaired t-test ****P<0.0001. *(E)* Immunofluorescence of HLA-G (green) in control shRNA and NOTUM shRNA treated cells following exposure to conditions that promote EVT cell differentiation. DAPI positive nuclei are shown in blue. Merged fluorescence images of HLA-G and DAPI (Scale bar: 500 μm). *(F and G)* A volcano plot and a heat map depicting RNA-sequencing analysis of control shRNA and NOTUM shRNA treated EVT cells (n=3 per group). *(F)* Blue dots represent significantly downregulated transcripts (P≤0.05) and a logarithm of base two-fold change of less than or equal to −2. Red dots represent significantly upregulated transcripts with P≤0.05 and a logarithm of base two-fold change of ≥2. *(G)* Heat map of control shRNA and NOTUM shRNA treated TS cells induced to differentiate into EVT cells. Colors represent z-scores of RPKM values. *(H)* Matrigel invasion assay of control shRNA or NOTUM shRNA treated cells under conditions promoting EVT cell differentiation. *(I)* The relative number of invading cells of control shRNA and NOTUM shRNA treated cells. Graphs represent mean values ± SEM, n=3, unpaired t-test, ****P<0.0001.

### Upstream regulation of NOTUM

We next sought to investigate the control of NOTUM expression in differentiating EVT cells. Inspection of RNA-seq profiles from earlier efforts of our laboratory indicated that endothelial PAS domain protein 1 (**EPAS1**) was a potential upstream regulator of NOTUM (**35**). EPAS1 protein has been referred to as hypoxia-inducible factor 2 alpha (**HIF2A**). NOTUM and EPAS1 exhibited parallel patterns of expression in first trimester human placenta and TS cells in the stem state and following their differentiation into EVT cells (**Fig. 5A-C**). Inhibition of EPAS1 expression in human TS cells resulted in the downregulation of NOTUM transcript and protein expression (**Fig. 5 D, E**). These observations support a role for EPAS1/HIF2A as an upstream regulator of *NOTUM*.

**Fig. 5.**
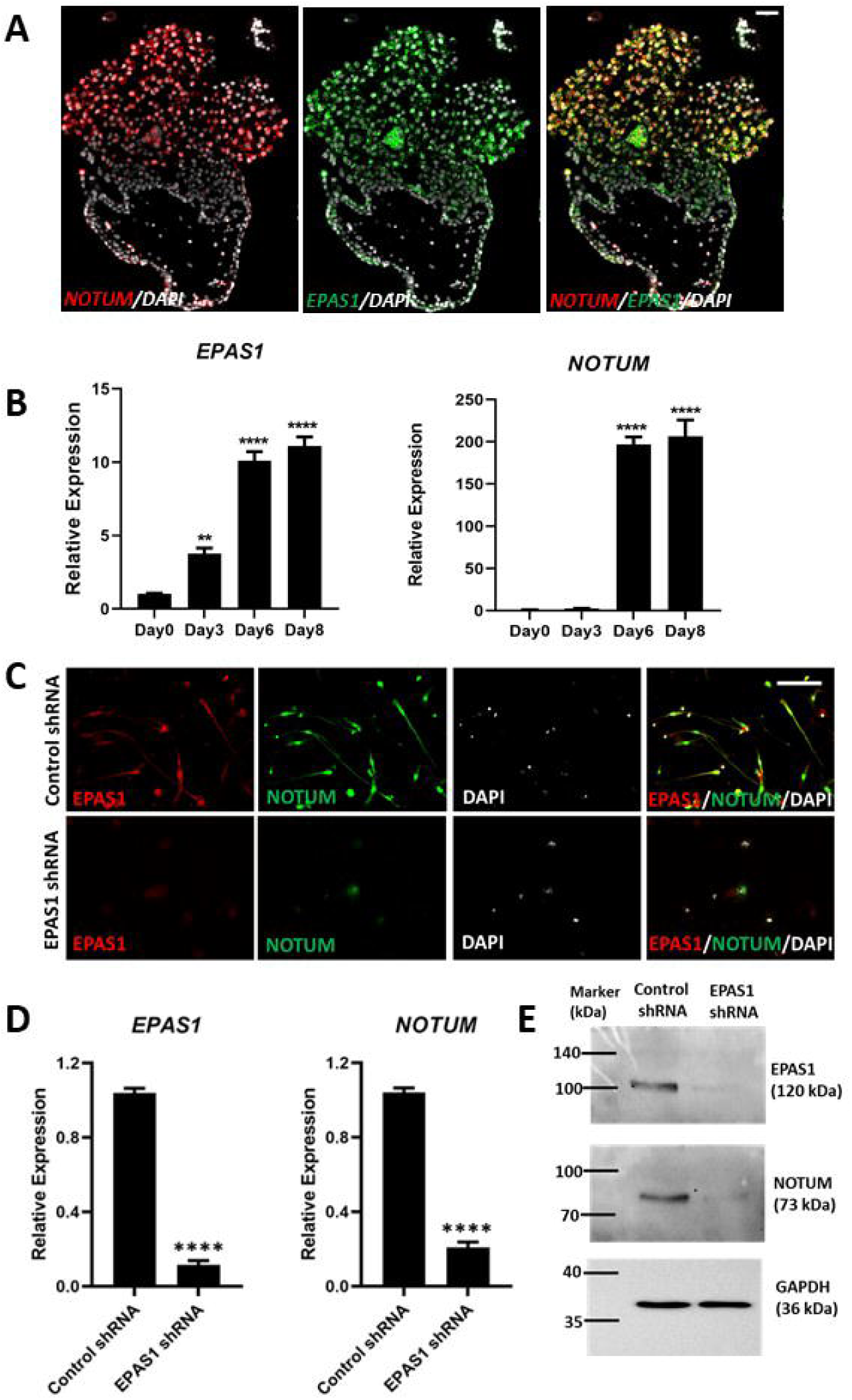
Upstream regulation of NOTUM in EVT cells. *(A)* In situ hybridization of first trimester human placenta at 12 weeks of gestation probed for *NOTUM* (red) and *EPAS1* (green). *NOTUM* (red) and *EPAS1* (green) are co-localized and expressed in the distal region of the EVT cell column but not in basal cytotrophoblast. DAPI positive nuclei are shown in grey (Scale bars: 50 μm). The next series of experiments were performed with X,X CT-27 cells. (*B*) Time course of *EPAS1* and *NOTUM* from stem state (Day 0) and days 3, 6, and 8 of EVT cell differentiation. Graphs represent mean values ± SEM, one-way analysis of variance, Tukey’s post hoc test, n=4, **P<0.01, ****P<0.0001. *(C)* Immunofluorescence of EPAS1/HIF2A (red) and NOTUM (green) in TS cells induced to differentiate into EVT cells following transduction with lentivirus containing control shRNA or EPAS1 shRNA. DAPI positive nuclei are shown in grey. Merged fluorescence images EPAS1, NOTUM and DAPI (Scale bar: 500 μm). Effects of EPAS1 shRNA knockdown on EPAS1 and NOTUM expression following 8 days of EVT cell differentiation assessed by RT-qPCR *(D*), graphs represent mean values ± SEM, n=3, unpaired *t*-test ****P<0.0001, and western blotting (*E*). GAPDH was used as a loading control for the western blot.

Collectively, the findings indicate that NOTUM is required for acquisition of structural, molecular, and functional indices of EVT cell differentiation and are consistent with its restraining actions on canonical WNT signaling.

## DISCUSSION

Our findings indicate that suppression of canonical WNT signaling is critical for human TS cell differentiation into EVT cells. In vitro EVT cell differentiation is accompanied by extensive cell elongation and spreading and the upregulation of transcripts indicative of the EVT cell fate (e.g., *HLA-G*, *MMP2*) (**25**). Consistent with these observations, we found that addition of a potent WNT activator, CHIR99021 (GSK3B inhibitor), inhibited differentiation of TS cells into EVT cells. We next interrogated human TS cells in stem and EVT differentiation states for expression of components of the WNT signaling pathway with the goal of identifying potential endogenous regulators of WNT signaling. Among the upregulated transcripts, NOTUM expression was striking in terms of both magnitude of the increase and overall expression level. NOTUM antagonizes WNT signaling by facilitating the depalmitoloylation of WNT proteins, which impairs their binding to frizzled receptors (**40**). Importantly, the differentiation-associated increase in NOTUM expression was required for optimal EVT cell differentiation. Upstream activation of NOTUM was regulated, at least in part, by the transcription factor, EPAS1. Glial cell missing 1 (**GCM1**) has also been identified as a potential upstream regulator of *NOTUM* (**41, 42**). EPAS1 may modulate *NOTUM* expression directly at the *NOTUM* locus or indirectly through stimulating *GCM1* expression (**35**). Collectively, our findings indicate that WNT drives trophoblast cell stemness, while EVT cell differentiation is facilitated by WNT suppression through the actions of NOTUM. In contrast to earlier reports (**29-31**), we did not find convincing evidence supporting a role for WNT signaling as a driver of EVT cell differentiation. There are important caveats associated with this statement. We did observe that EVT cell behavior within the decidual bed exhibited heterogeneity regarding WNT signaling. Thus, WNT promotion of EVT cell differentiation may be context dependent and not readily demonstrated with human TS cells cultured as described in this report. Additionally, it is important to appreciate that NOTUM is not a binary switch (“on” versus “off”) controlling EVT cell differentiation but instead likely acts as a rheostat to modulate WNT signaling. Too much activation via the WNT signaling pathway appears incompatible with EVT cell differentiation; however, this does not negate that some WNT signaling may be required for EVT cell differentiation.

Despite the recognized significance of WNT signaling in trophoblast cell biology (**43-45**), a regulatory role for NOTUM in this process has not been recognized. Nonetheless, NOTUM dysregulation is associated with placenta pathology and pregnancy related disorders (**46-49**). WNT signaling is a conserved regulator of placentation in the human and mouse (**43-45**). In contrast, NOTUM appears to be a species-specific regulator of the invasive trophoblast cell lineage, which may be best exemplified in comparing rat with human placentation. The rat exhibits robust intrauterine trophoblast cell invasion (**50, 51**) but invasive trophoblast cells do not express detectable NOTUM, instead rat invasive trophoblast cells express other WNT signaling antagonists, including catenin beta interacting protein 1 (**CTNNBIP1**) and NKD inhibitor of WNT signaling pathway 1 (**NKD1**) (**52**). It remains to be determined whether and how WNT signaling and WNT antagonists may modulate development and function of the rat invasive trophoblast cell lineage.

Overall, canonical WNT signaling is pivotal to the maintenance of human TS cell stemness and regulation of EVT cell differentiation, especially within the EVT cell column. NOTUM is identified as a critical liberator from canonical WNT dominance leading to development of EVT cells. Thus, the appropriate modulation of canonical WNT signaling is critical to placental development and its dysregulation a potential factor in placental disease. As EVT cells move into the uterine decidua a subset of EVT cells acquire responsiveness to WNT signaling and their dependence on NOTUM subsides.

## MATERIALS AND METHODS

### Human placentation site specimens

Sections from paraffin-embedded deidentified first trimester human placenta and placental bed tissue specimens were obtained from the Lunenfeld-Tanenbaum Research Institute (Mount Sinai Hospital, Toronto, Canada) and St. Mary’s Hospital, Manchester, United Kingdom, respectively. Tissue collections were performed after written informed consent. Institutional approval was obtained from Human Research Ethics Review Committees at the University of Toronto, Central Manchester Health Trust, and the University of Kansas Medical Center (**KUMC**).

### TS cell culture

Cytotrophoblast-derived CT27 (X,X) and CT29 (X,Y) lines were used in the experiments. Human TS cells were maintained in stem state conditions or induced to differentiate into syncytiotrophoblast or EVT cells as previously described (**25**). Human TS cells were cultured in six-well plates precoated with 5 μg/mL of Corning^TM^ collagen IV (CB40233, Thermo Fisher) and Basal Human TS Cell Medium [Dulbecco’s Modified Eagle Medium/F12 (DMEM/F12, 11320033, Thermo Fisher) containing 100 μM 2-mercaptoethanol, 0.2% (vol/vol), fetal bovine serum (**FBS**), 50 μM penicillin, 50 U/mL streptomycin, 0.3% bovine serum albumin (BP9704100, Thermo Fisher), 1% Insulin-Transferrin-Selenium-Ethanolamine solution (**ITS-X**, vol/vol, Thermo Fisher), supplemented with 1.5 μg/mL L-ascorbic acid (A8960, Sigma-Aldrich), 50 ng/mL epidermal growth factor (**EGF**, E9644, Sigma-Aldrich), 2 μM CHIR99021 (04-0004, Reprocell), 0.5 μM A83-01 (04-0014, Reprocell), 1 μM SB431542 (04-0010, Reprocell), 0.8 mM valproic acid (P4543, Sigma-Aldrich), and 5 μM Y27632 (04-0012-02, Reprocell).

EVT cell differentiation was performed as previously described (**25**). Human TS cells were plated onto a 6-well plate precoated with 1 μg/mL collagen IV at a density of 1 × 10^5^ cells per well and cultured in EVT Cell Differentiation Medium, which consists of Basal Human TS Cell Medium supplemented with 100 ng/mL of neuregulin 1 (**NRG1**, 5218SC, Cell Signaling), 7.5 μM A83-01, 2.5 μM Y27632, 4% KnockOut Serum Replacement (**KSR**, 10828028, Thermo Fisher), and 2% Matrigel (CB-40234, Fisher). On day 3 of EVT cell differentiation, the medium was replaced with the EVT Cell Differentiation Medium without NRG1, and the Matrigel^®^ concentration was decreased to 0.5%. On day 6 of EVT cell differentiation, the medium was replaced with EVT Cell Differentiation Medium without NRG1 or KSR and with Matrigel^®^ at a concentration of 0.5%. Cells were analyzed on day 8 of EVT cell differentiation or in one set of experiments, the culture was extended to day 14 of EVT cell differentiation.

TS cells were induced to differentiate into syncytiotrophoblast using three-dimensional (3D) culture conditions [**ST(3D)**] (**25**). TS cells (2.5 x 10^5^) were seeded into six cm Petri dishes and cultured in 3 mL of ST(3D) medium containing DMEM/F12 supplemented with 100 μM 2-mercaptoethanol, 0.5% penicillin-streptomycin, 0.3% BSA, 1% ITS-X supplement, 2.5 μM Y27632, 50 ng/ml EGF, 2 μM forskolin (F6886, Sigma-Aldrich), and 4% KSR. An equal amount of fresh ST(3D) media was added at day 3. Cells were passed through a 40 μm mesh filter to remove dead cells and debris at day 6. Cells remaining on the 40 μm mesh filter were collected and analyzed.

### shRNAs, transient transfection, lentivirus production and transduction

A lentiviral-mediated shRNA delivery approach was used to silence *NOTUM* and *EPAS1* gene expression. *NOTUM* shRNAs were designed and subcloned into the pLKO.1 vector at *AgeI* and *EcoRI* restriction sites. *EPAS1* shRNAs were described previously (**35**). shRNA sequences used in the analyses are included in **Table S1**. Lentiviral packaging vectors were obtained from Addgene and included pMDLg/pRRE (plasmid 12251), pRSVRev (plasmid 12253), and pMD2.G (plasmid 12259). Lentiviral particles were produced following transient transfection of the shRNA-pLKO.1 vector and packaging plasmids into Lenti-X cells (632180; Takara Bio USA) using Attractene (301005; Qiagen) in Opti-MEM I (51985-034; Thermo Fisher). Thereafter, cells were maintained in DMEM (11995-065; Thermo Fisher) with 10% FBS. Culture supernatants containing lentiviral particles were collected every 24 h for two days and stored frozen at -80°C until use.

Human TS cells were plated at 80,000 cells per well in 6-well tissue culture-treated plates coated with 5 μg/mL collagen IV (CB40233; Thermo Fisher) and incubated for 24 h. Just before transduction, medium was changed, and cells were incubated with 2.5 μg/mL polybrene for 30 min at 37°C. Immediately following polybrene incubation, TS cells were transduced with 500 μL of lentiviral particle supernatant and then incubated for 24 h. Medium was changed at 24 h post-transduction and selected with puromycin dihydrochloride (5 μg/mL, A11138-03; Thermo Fisher) for 2 days. Cells were allowed to recover for 1 to 3 days in complete human TS medium before splitting for EVT cell differentiation.

### Matrigel invasion assay

Invasiveness was assessed by plating cells on Matrigel-coated transwell inserts with 8.0 μm transparent polyester membrane pores (Corning, 353097), as previously described (**53**). Briefly, EVT cells were dissociated into single cells and seeded at a density of 2 × 10^5^ cells per well into the upper chamber of Matrigel-coated transwells in 200 μL EVT Cell Differentiation Medium. The lower chamber was filled with 750 μL of the EVT Cell Differentiation Medium containing 20% FBS. EVTs were cultured at 37°C in 5% CO_2_. After 36 h, a cotton swab moistened with medium was inserted into the top of the Matrigel matrix-coated permeable support (apical side) and gentle but firm pressure was used to remove cells from the support. The lower chamber was fixed with 4% paraformaldehyde, washed with phosphate buffered saline (pH 7.4), and next stained with crystal violet. Invaded cells were imaged on a Nikon Eclipse 80i microscope. Subsequently, stained cells from three random fields were counted to calculate the relative fold change in the number of invading cells.

### Immunofluorescence

Human TS cells or differentiated EVT cells were fixed with 4% paraformaldehyde (Sigma-Aldrich) for 20 min at room temperature. Immunofluorescence analysis was performed using a primary antibody against NOTUM (1:500, SAB3500082, Sigma-Aldrich), active-CTNNB1 (1:400, clone 8E7, 05-665, Sigma-Aldrich), HLA-G (1:400, ab52455; Abcam), or EPAS1/HIF2A (1:500, D6T8V, rabbit monoclonal antibody 59973, Cell Signaling). Alexa Fluor 488 goat anti-mouse IgG (1:800, A32723 Thermo Fisher), Alexa Fluor 568 goat anti-mouse IgG (1:800, A11031, Thermo Fisher), Alexa Fluor 568 goat anti-rabbit IgG, (1:800, A10042, Thermo Fisher) were used to detect locations of the primary antibody-antigen complexes within cells and tissues. For immunohistochemical analysis, paraffin-embedded slides incubated with 10% normal goat serum (50062Z; Thermo Fisher Scientific) for 1 h. Sections were incubated overnight with primary antibodies: NOTUM (1:500, SAB3500082, Sigma-Aldrich), active-CTNNB1 (1:400, clone 8E7, 05-665, Sigma-Aldrich or 1:400, clone 8814, D13A1, Cell Signaling Technology), or HLA-G (1:400, ab52455; Abcam). After washing with phosphate-buffered saline (pH 7.4), sections were incubated for 2 h with corresponding secondary antibodies: Alexa Fluor 488 goat anti-mouse IgG (1:800, A32723 Thermo Fisher), Alexa Fluor 568 goat anti-mouse IgG (1:800, A11031, Thermo Fisher), Alexa Fluor 568 goat anti-rabbit IgG, (1:800, A10042, Thermo Fisher). Nuclei were visualized by staining with 4’6’-diamidino-2-phenylindole (**DAPI**, Molecular Probes). Immunostained sections were mounted in Fluoromount-G (0100-01; SouthernBiotech). Images were captured on a Nikon 90i upright microscope with a Roper Photometrics CoolSNAP-ES monochrome camera and 2047 Zeiss Axio Observer 7 with Apotome III and Artificial Intelligence (**AI**) Sample Finder.

### Western blot analysis

Cell lysates were prepared by sonication in radioimmunoprecipitation assay lysis buffer (sc-24948A, Santa Cruz Biotechnology) supplemented with Halt protease and phosphatase inhibitor mixture (78443, Thermo Fisher). Nuclear and cytosolic fraction isolated from subcellular protein fractionation kit (78840, Thermo Fisher) according to the manufacturer’s instructions. Protein concentrations were measured using the DC Protein Assay (5000113-115, Bio-Rad). Proteins (20 μg/lane) were separated by sodium dodecyl sulfate polyacrylamide gel electrophoresis and transferred onto polyvinylidene difluoride membranes (10600023, GE Healthcare). After transfer, membranes were blocked with 5% non-fat milk/BSA in Tris buffered saline with 0.1% Tween 20 (**TBST**) and probed with primary antibodies to NOTUM (1:1500, SAB3500082, Sigma-Aldrich), HLA-G (1:1000. ab52455, Abcam), total CTNNB1 (1:1000, clone D10A8, 8480, Cell Signaling), active-CTNNB1 (1:1000, clone 8E7, 05-665, Sigma-Aldrich), active-CTNNB1 (1:500, 8814 Cell Signaling), phosphorylated CTNNB1 (1:1000, Ser33/37/Thr41, 9561 Cell Signaling), EPAS1 (1:300, 66731-1-Ig, Proteintech) and glyceraldehyde-3-phosphate dehydrogenase (ab9485; Abcam) overnight at 4°C. Membranes were washed three times for five min each with TBST and then incubated with secondary antibodies (goat anti-rabbit IgG HRP, A0545, Sigma Aldrich; goat anti-mouse IgG HRP, 7076; Cell Signaling) for 1 h at room temperature. Immunoreactive proteins were visualized by enhanced chemiluminescence according to the manufacturer’s instructions (Amersham).

### In situ hybridization

Detection of NOTUM transcripts was performed on paraffin-embedded human placentation site tissue sections using the RNAscope Multiplex Fluorescent Reagent Kit version 2 (Advanced Cell Diagnostics), according to the manufacturer’s instructions. RNAscope probes were used to detect *NOTUM* (NM_178493.5, 430311, target region: 259-814), *PLAC8* (NM_016619.3, 858491-C2, target region: 5-1448), *EPAS1* (NM_001430.4, 41059-C1, target region:1332 - 2354), and *CDH1* (NM_004360.3, 311091-C2, target region: 263-1255). Fluorescence images were captured on a Nikon 90i upright microscope (Nikon) with a Roper Photometrics CoolSNAP-ES monochrome camera (Roper) and 2047 Zeiss Axio Observer 7 with Apotome III and AI Sample Finder.

### RNA isolation, cDNA synthesis, and RT-qPCR

Total RNA was isolated from cells with TRIzol reagent (15596018, Thermo Fisher) as described previously (**54**). cDNA was synthesized from 1 μg of total RNA using a High-Capacity cDNA Reverse Transcription Kit (4368813; Thermo Fisher) and diluted 10 times with Milli-Q water. Reverse transcriptase-polymerase chain reaction (**RT-qPCR**) was performed using a reaction mixture containing PowerSYBR Green PCR Master Mix (4367659, Thermo Fisher) and primers (250 nM each). PCR primer sequences are presented in **Table S2**. Real-time PCR amplification and fluorescence detection were carried out using a QuantStudio 7 Flex Real-Time PCR System (Thermo Fisher). An initial step (95 °C, 10 min) preceded 40 cycles of a two-step PCR at the following conditions: 92 °C, for 15 s and 60 °C for 1 min, followed by a dissociation step (95 °C for 15 s, 60 °C for 15 s, and 95 °C for 15 s). The comparative cycle threshold method was used for relative quantification of the amount of mRNA for each sample normalized to a housekeeping gene *GAPDH*.

### RNA-seq analysis

RNA-seq was performed as previously described (**55**). Transcript profiles were generated from vehicle control and CHIR99021 (2 μM) or control shRNA and NOTUM shRNA exposures in CT27 human TS cells cultured in conditions promoting EVT cell differentiation (n=3 experiments per group, each sample corresponded to a distinct transduction). Complementary DNA libraries from total RNA samples were prepared with Illumina TruSeq RNA preparation kits according to the manufacturer’s instructions. RNA integrity was assessed using an Agilent 2100 Bioanalyzer. Barcoded cDNA libraries were multiplexed onto a TruSeq paired-end flow cell and sequenced (100-bp paired-end reads) with a TruSeq 200-cycle SBS kit (Illumina). Samples were run on Illumina HiSeq2500 sequencers located at the KUMC Genome Sequencing Facility. Reads from *.fastq files were mapped to the human reference genome (GRCh37) using CLC Genomics Workbench 12.0 (Qiagen). Transcript abundance was expressed as reads per kilobase of transcript per million mapped reads (**RPKM**), DEGs were identified using false discovery rate (**FDR**) and fold change (**FC**) versus control groups (FDR<0.05; FC≥1.5). Statistical significance was calculated by empirical analysis of digital gene expression followed by Bonferroni’s correction. QIAGEN Ingenuity Pathway Analysis of DEGs was performed to gain additional insight into the biological impacts of the cell/molecular manipulations.

### Statistical analysis

Statistical analyses were performed with GraphPad Prism 9 software. Student’s *t* test, one-way or two-way analysis of variance were applied as appropriate. Statistical significance was determined as P<0.05. All values are presented as the mean ± standard error of the mean (**SEM**) of at least three independent experiments.

## Data availability

The datasets generated and analyzed for this study have been deposited in the Gene Expression Omnibus (GEO) database, https://www.ncbi.nlm.nih.gov/geo/ (accession no. GSE206036).

## ACKNOWLEDGMENTS

The research was supported by postdoctoral fellowships by the KUMC Biomedical Research Training Program and K-INBRE (P20 GM103418; VS, AM, MK), a Lalor Foundation Fellowship (AM), and an NIH National Research Service Award postdoctoral fellowship (HD096809, KMV) and NIH grants (HD020676, HD079363, HD099638, HD105734), and the Sosland Foundation. We thank Stacy Oxley and Brandi Miller for administrative assistance.

