## Supplemental File for "NOTUM-MEDIATED WNT SILENCING DRIVES EXTRAVILLOUS TROPHOBLAST CELL LINEAGE DEVELOPMENT"

**This PDF file includes:**

Figures S1 to S10

Tables S1 and S2

**Other supporting materials for this manuscript include the following:**

Datasets1 to 3

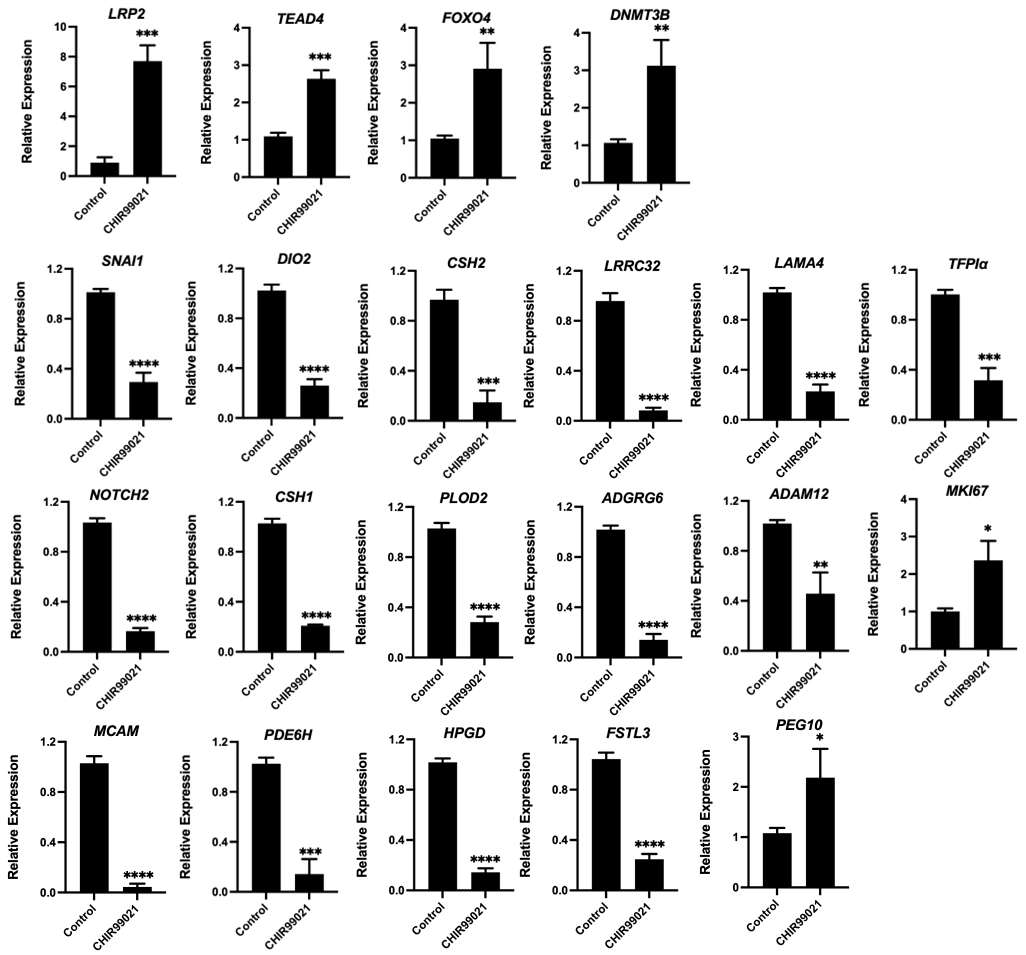

**Fig. S1.** RT-qPCR validation of selected differentially expressed transcripts obtained from RNA-sequencing of vehicle or CHIR99021 (2 μM) treated X,X CT-27 EVT cells following 8 days of EVT cell differentiation (n = 3). Graphs represent mean values ± SEM, unpaired t test, ****P<0.0001, ***P<0.001, **P<0.01, *P<0.05).

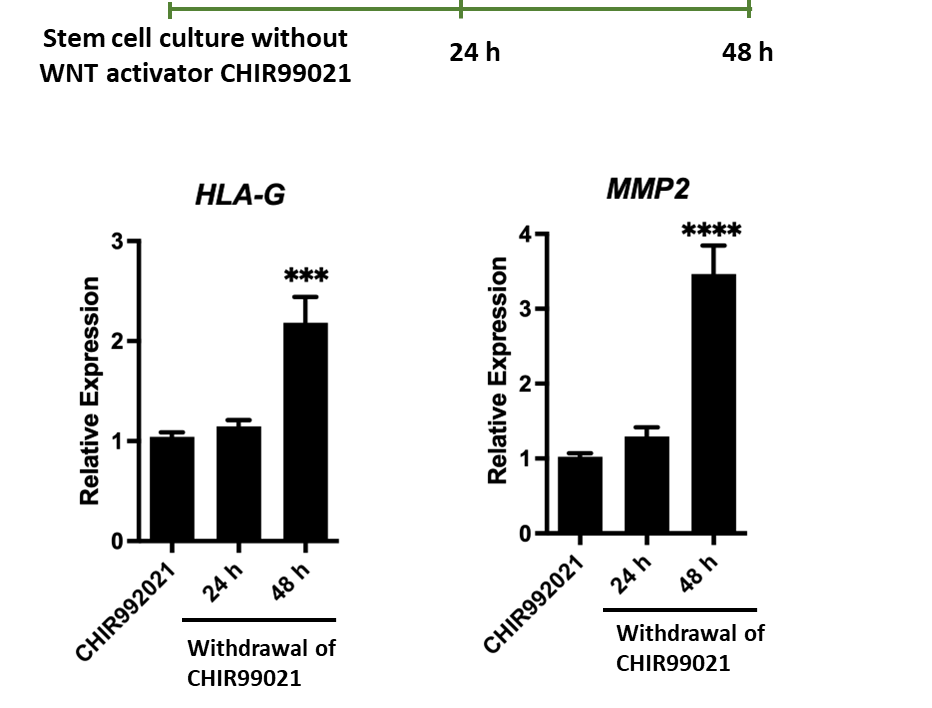

**Fig. S2.** Effect of removal of CHIR99021 in X,X CT-27 TS cells maintained in the stem state on the expression of EVT cell associated transcripts (*HLA-G* and *MMP2*). Transcript levels of *HLA-G* and *MMP2* increased within 48 h of CHIR99021 withdrawal (n=3). Graphs represent mean values ± SEM, one-way analysis of variance, Tukey’s post hoc test, ****P<0.0001, ***P<0.001.

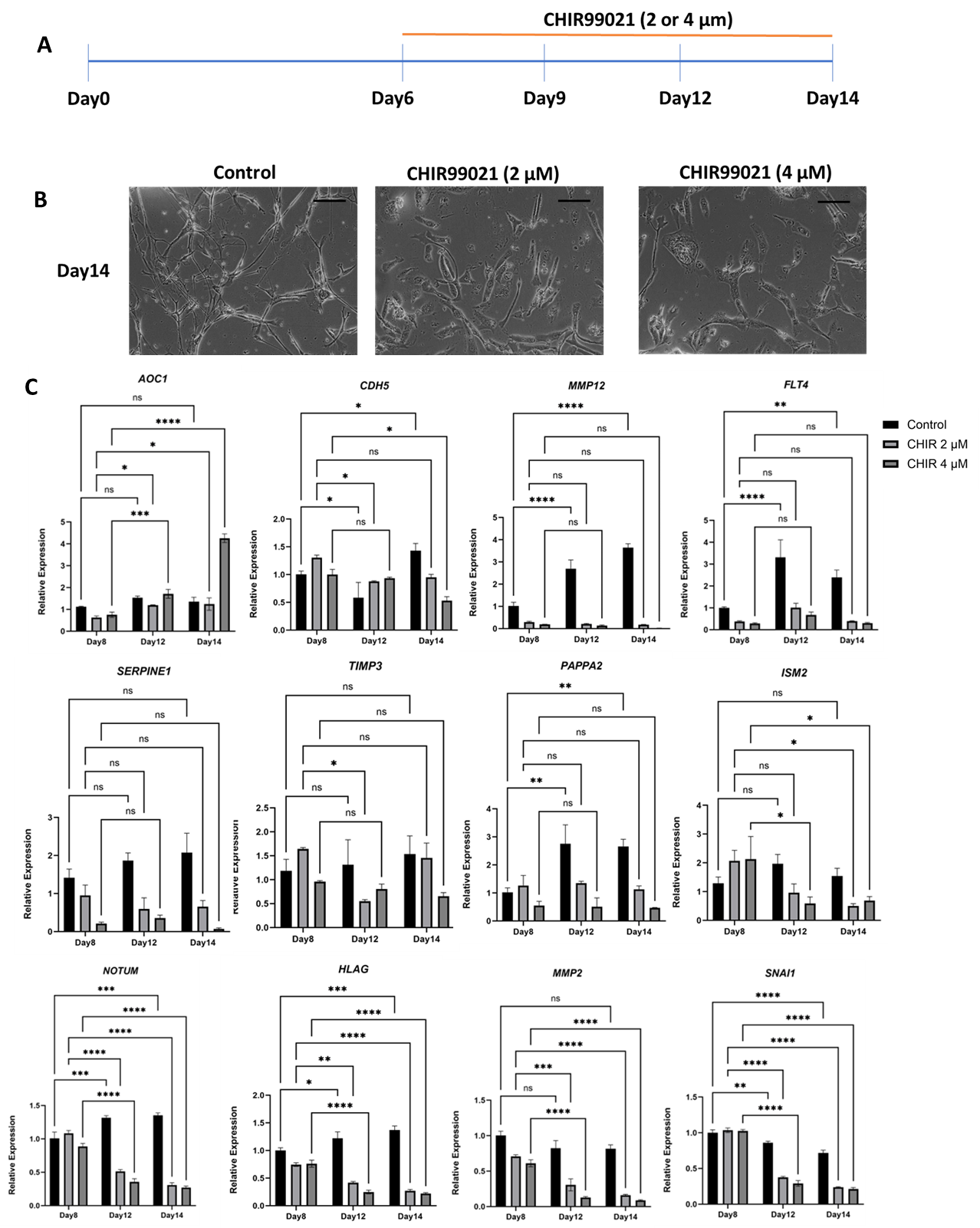

**Fig. S3.** EVT cell associated transcript expression following exposure to CHIR99021 (WNT activation, 2 or 4 μM) during extended X,X CT-27 EVT cell culture (n=3). Graphs represent mean values ± SEM, two-way analysis of variance, ****P<0.0001, ***P<0.001, **P<0.01, *P<0.05.

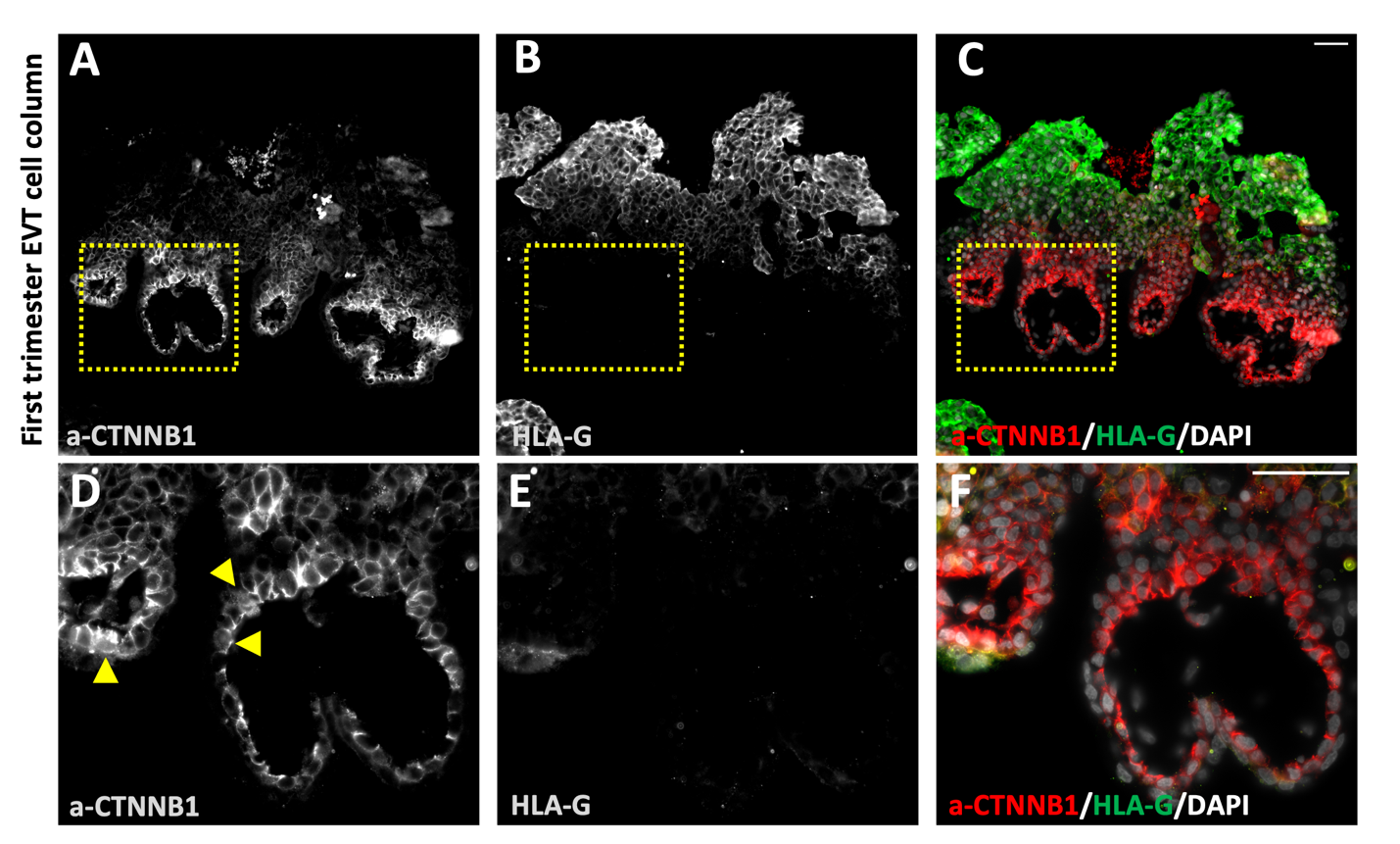

**Fig. S4.** Immunofluorescence of first trimester human placenta at 6 weeks of gestation tested for the presence of active CTNNB1 (**a-CTNNB1**) and HLA-G**.** *(A)* a-CTNNB1 is expressed in cytotrophoblast. *(B)* HLA-G is expressed in EVT column cells but not in cytotrophoblast or syncytiotrophoblast compartments**.** *(C)* Merged fluorescence images of a-CTNNB1 (red) and HLA-G (green). (D-F) Higher magnification images of yellow inset shown in panels A-C, respectfully. Yellow arrowheads indicate a-CTTNB1 in cytotrophoblast nuclei. Scale bar: 50 μm.

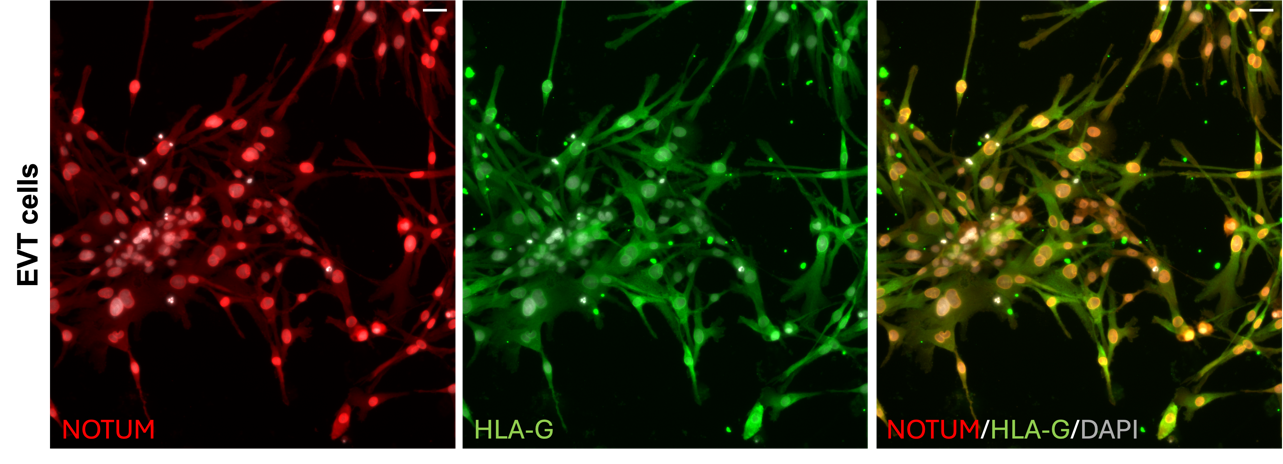

**Fig. S5.** Immunolocalization of NOTUM (red) and HLA-G (green) in EVT cells**.** Merged image of NOTUM, HLA-G, and DAPI immunofluorescence is shown in the right panel (Scale bars: 50 μm).

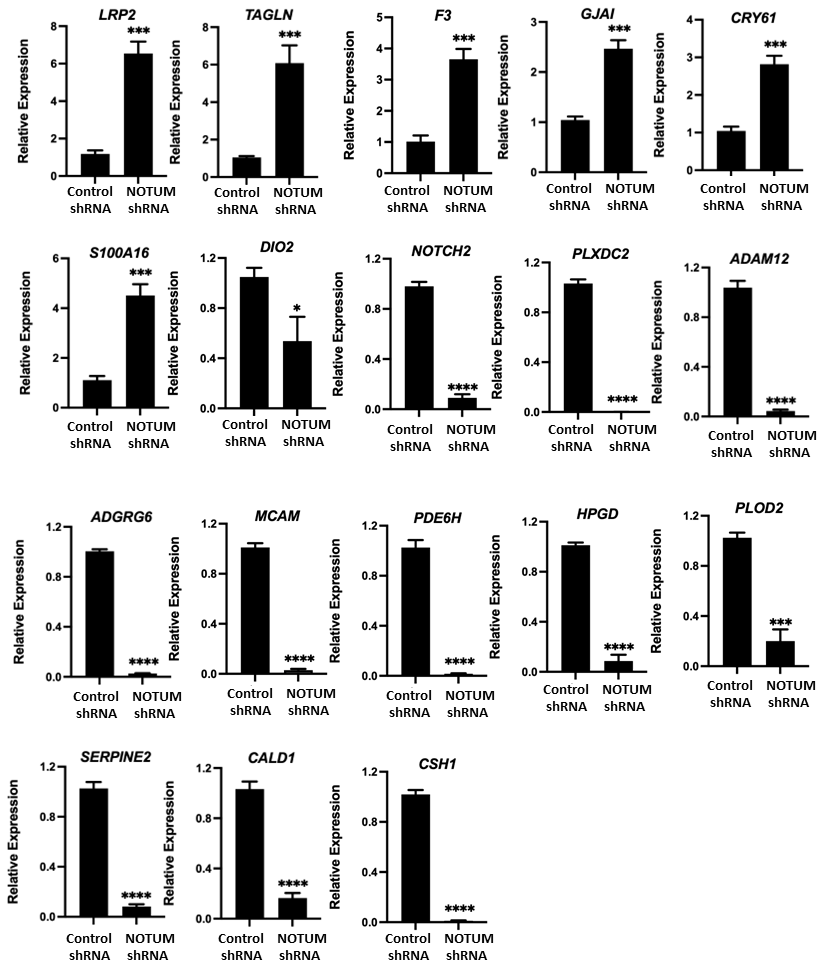

**Fig. S6.** RT-qPCR validation of selected differentially expressed transcripts obtained from RNA-sequencing of control shRNA or NOTUM shRNA treated X,X CT-27 following 8 days of EVT cell differentiation (n = 3). Graphs represent mean values ± SEM, unpaired t test, ****P<0.0001, ***P<0.001, **P<0.01.

**Fig. S7.** Effect of NOTUM inhibition on canonical WNT signaling. Experiments were performed with X,X CT-27 cells following 8 days of EVT cell differentiation. *(A)* Immunolocalization of NOTUM (green) and active CTNNB1 (**a-CTNNB1**; red) protein in control shRNA or NOTUM shRNA treated EVT cells**.** DAPI positive nuclei are shown in blue. Merged images of NOTUM, a-CTNNB1, and DAPI immunofluorescence (Scale bars: 500 μm). *(B)* Western blot images of CTNNB1, a-CTNNB1 and phospho-CTNNB1 (Ser33/37/Thr41) in cytosolic and nuclear enriched fractions of control shRNA or NOTUM-specific shRNA treated EVT cells.

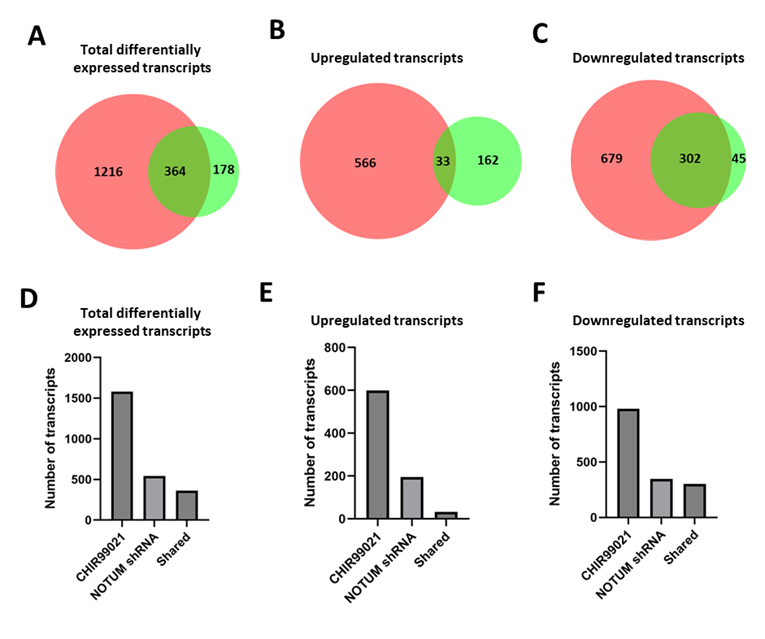

**Fig. S8.** Comparison of RNA sequencing datasets from CHIR9021 and NOTUM shRNA exposures (see Datasets 1 and 2). Venn diagrams showing total differentially expressed transcripts (*A)*, upregulated transcripts (*B*), and downregulated transcripts (*C*). The numbers of transcripts shared in both datasets are also presented. The same data is presented in bar graph format (*D-F*). Shared upregulated and downregulated transcript lists are provided in Dataset 3.

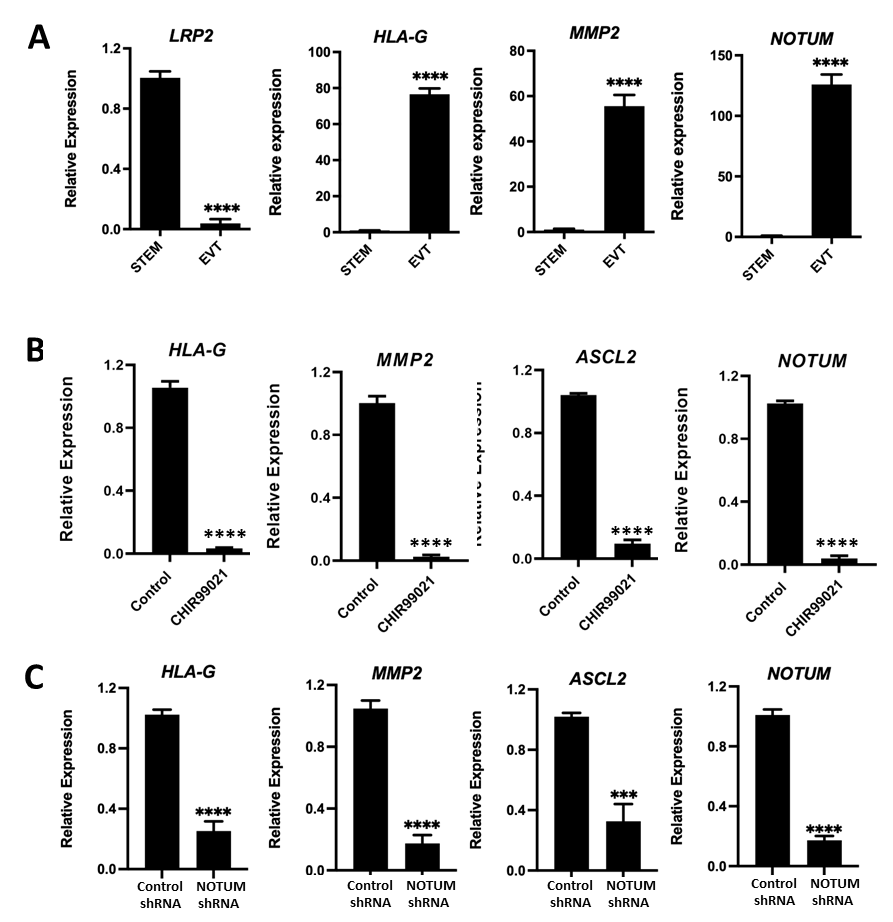

**Fig. S9.** WNT signaling antagonizes and NOTUM is required for EVT cell differentiation in X,Y CT-29 human TS cells in the stem state or following 8 days of EVT cell differentiation. *(A)* LRP2 transcript is decreased and *HLA-G*, *MMP2*, *NOTUM* transcripts are increased as TS cells transition from the stem state to EVT cells, ****P<0.0001. *(B)* Transcript levels of four invasive trophoblast cell markers (*HLA-G, MMP2, ASCL2* and *NOTUM*) on day 8 of EVT differentiation following exposure to vehicle or CHIR99021 (2 μM), ****P<0.0001. *(C)* Transcript levels of four EVT cell markers (*HLA-G, MMP2, ASCL2* and *NOTUM*) on day 8 of EVT differentiation following transduction with lentivirus containing a control shRNA or NOTUM shRNA. NOTUM depletion decreased levels of transcripts associated with the EVT cell phenotype. The results are presented as mean values ± SEM, n=3, unpaired t-test, ****P<0.0001. All transcript levels were measured by RT-qPCR.

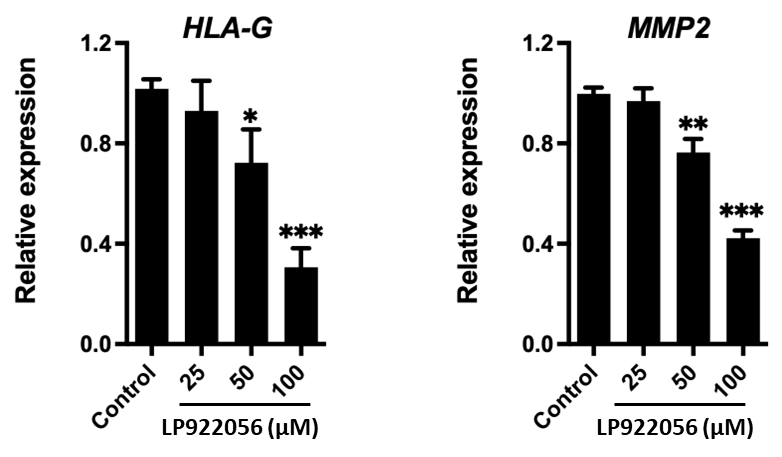

**Fig. S10.** Effects of LP922056 (small molecule inhibitor of NOTUM) on X,X CT-27 EVT cell differentiation. Cells were treated with vehicle or various concentrations of LP922056 (0-100 μM) during EVT cell differentiation. On day 8 of EVT cell differentiation, cells were harvested and transcripts for *HLA-G* and *MMP2* measured by RT-qPCR (n=3). Graphs represent mean values ± SEM, one-way analysis of variance, Tukey’s post hoc test, ***P<0.001, **P<0.01, *P<0.05.

**Table S1. shRNA sequences**

| **shRNA** | **Sequences** |
| --- | --- |
| NOTUM shRNA | AACTGCTTCTTTGGCTACAA |
| EPAS1 shRNA | GCGCTCAGCCGTGGAGTACAT |

**Table S2. Primer sequences for qRT-PCR analyses**

| **Target** | **Species** | **Forward** | **Reverse** |
| --- | --- | --- | --- |
| *HLA-G* | Human | CCACCACCCTGTCTTTGACTAT | ACGTCCTGGGTCTGGTCCT |
| *MMP2* | Human | TGGCACCCATTTACACCTACAC | ATGTCAGGAGAGGCCCCATAGA |
| *ASCL2* | Human | GCACCAACACTTGGAGATTTT | AATGGATTCTCTGTGCCCTTAG |
| *NOTUM* | Human | ACAGGGATCCTGTCCTCACA | CTCCAAACATCACTGGAGCA |
| *LRP2* | Human | CTGCTCCTGGCTCTCGTC | TCCCATCACACCTCCAGTCT |
| *TEAD4* | Human | CAGGTGGTGGAGAAAGTTGAGA | GTGCTTGAGCTTGTGGATGAAG |
| *FOXO4* | Human | AGCAACAGCTCAGCAGGATG | TGGCCTCGTTGTGAACCTTG |
| *DNMT3B* | Human | AAGTCGAAGGTGCGTCGTG | GAAGCCATTTGTTCTCGGCTC |
| *SNAI1* | Human | TACAGCGAGCTGCAGGACT | ATCTCCGGAGGTGGGATG |
| *DIO2* | Human | GGTTCCCCTTCACCCTCTCTG | TCTCTGCCACAGTCTCATAGGT |
| *CSH2* | Human | GCTCACCTAGCGGCAATG | CCGTCTTCCAGCCTAAACTCCT |
| *LRRC32* | Human | CTGTGAGCTAGTCTGCCCTG | GGGTCTCATGGCTCCAAAGT |
| *LAMA4* | Human | GCCAAGAACTGTGCAGTGTG | CAGACGCACTTATCACAGCC |
| *TFPIa* | Human | ATGGAACCCAGCTCAATGCT | GGCACGACACAATCCTCTGT |
| *NOTCH2* | Human | AAGGAACCTGCTTTGATGACA | CAGGGAGCCAATACTGTCTGA |
| *CSH1* | Human | CATGACTCCCAGACCTCCTTCT | ATTTCTGTTGCGTTTCCTCCAT |
| *PLOD2* | Human | GCGTTCTCTTCGTCCTCATC | CATGAAGCTCCAGCCTTTTC |
| *ADGRG6* | Human | CTTCCACTGTGCTATGAAGGAG | TGGTAGCTGTCTTACTCCAATCTG |
| *MCAM* | Human | GGGTACCCCATTCCTCAAGT | CAGTCTGGGACGACTGAATG |
| *PDE6H* | Human | TGGAGGGGCTAGGAACAGA | GAGCGAGCTCATGCAATTC |
| *HPGD* | Human | CAGAAGACTCTGTTCATCCAGTG | TGTCCAGTCTTCCAAAGTGGT |
| *FSTL3* | Human | CTACATCTCCTCGTGCCACA | TCTTCTGCAGACTCACCACCT |
| *ADAM12* | Human | TGTGGAAGAAGGAGAGGAGTG | CATTGCAGCAGCGATTCATA |
| *PEG10* | Human | GGAGAACAGCGGAGAAGGTC | CAAAACCCGCTTATTTCACGC |
| *MKI67* | Human | TGACCCTGATGAGAAAGCTCAA | CCCTGAGCAACACTGTCTTTT |
| *F3* | Human | ACCGGGCTGTCTGTACTCTT | TTCCTAAGCCTCCGGGATGT |
| *GJAI* | Human | CTGAGTGCCTGAACTTGCCT | CTGGGCACCACTCTTTTG C |
| *CRY61* | Human | AAACCCGGATTTGTGAGGT | GCTGCATTTCTTGCCCTTT |
| *YAP1* | Human | GAACTCGGCTTCAGGTCCTC | GGTTCATGGCAAAACGAGGG |
| *TAGLN* | Human | CTGAGGAAGCCTTCTTTCCCC | GGGCCACACTGCACTATGAT |
| *S100A16* | Human | ACAGATTCTGGGAGTGCAGC | CTGGCGGGGCCTGCT |
| *PLXDC2* | Human | GGTAGACACGAACCGAGCAA | GGACAGATTCACTCTCTCGATCT |
| *CALD1* | Human | CTGTGCAGAAAAGTGGTGTCA | ACACCTTCAGCAGGAACAGG |
| *SERPINE2* | Human | TTGCAAAAATAACAACAGGGTCA | TGCAGTTGTTGCTGCTGAAG |
| *NCAM1* | Human | CTGGGAACTGCAGCAAAACC | TCCGGGTAGAAGTCCTCCAG |
| *PAPPA2* | Human | GAGCAAAACACTTGGAACCCA | TGATGCTTGGGGTTGTTGGT |
| *SERPINE1* | Human | CACAACCCCACAGGCCCTT | AGGGTGTTTCTTCCACTGGC |
| *FLT4* | Human | GTGACGTGTGGTCCTTTGG | CTGGCAGAACTCCTCATTGAT |
| *CDH5* | Human | ATGAGATCGTGGTGGAAGCG | AATGTGTACTTGGTCTGGGTGAA |
| *TIMP3* | Human | GCTGGAGGTCAACAAGTACCA | CACAGCCCCGTGTACATCT |
| *ISM2* | Human | CAGAATTGGTCCACGCAACC | GGCTCAGCCAACAGGTCTAT |
| *MMP12* | Human | TGGTTTTTGCCCGTGGAGCTCAT | GAATGGCCAATCTCGTGAACAGCA |
| *GAPDH* | Human | CCTCAACGACCACTTTGTCAAG | TCTTCCTCTTGTGCTCTTGCTG |

**Dataset S1 (separate file).** Differentially regulated transcripts identified by RNA-seq analysis in day 8 EVT differentiated human X,X CT-27 TS cells following CHIR99021 treatment.

**Dataset S2 (separate file).** Differentially regulated transcripts identified by RNA-seq analysis in day 8 EVT differentiated human X,X CT-27 TS cells following NOTUM knockdown.

**Dataset S3 (separate file).** Shared gene identified by RNA-seq analysis in EVT differentiated human X,X CT-27 TS cells following CHIR99021 treatment and NOTUM knockdown cells.
